# DeeplyEssential: A Deep Neural Network for Predicting Essential Genes in Microbes

**DOI:** 10.1101/607085

**Authors:** Md Abid Hasan, Stefano Lonardi

## Abstract

Essential genes are genes that critical for the survival of an organism. The prediction of essential genes in bacteria can provide targets for the design of novel antibiotic compounds or antimicrobial strategies. Here we propose a deep neural network (DNN) for predicting essential genes in microbes. Our DNN-based architecture called DeeplyEssential makes minimal assumptions about the input data (i.e., it only uses gene primary sequence and the corresponding protein sequence) to carry out the prediction, thus maximizing its practical application compared to existing predictors that require structural or topological features which might not be readily available. Our extensive experimental results show that DeeplyEssential outperforms existing classifiers that either employ down-sampling to balance the training set or use clustering to exclude multiple copies of orthologous genes. We also expose and study a hidden performance bias that affected previous classifiers.

The code of DeeplyEssential is freely available at https://github.com/ucrbioinfo/DeeplyEssential

## 1 Introduction

Essential genes are those genes that are critical for the survival and reproduction of an organism [17]. Since the disruption of essential genes induces the death of an organism, the identification of essential genes can provide targets for new antimicrobial/antibiotic drugs [7, 13]. The set of essential genes is also critical for the creation of artificial self-sustainable living cells with a minimal genome [16]. Essential genes have also been a cornerstone in understanding the origin and evolution of organisms [18].

The identification of essential genes via wet-lab experiments is labor intensive, expensive and time consuming. Such experimental procedures include single gene knock-out [3, 12], RNA interference and transposon mutagenesis [8, 32]. Moreover, these experimental approaches can produce contradicting results [23]. With the recent advances in high-throughput sequencing technology, computational methods for predicting essential genes has become a reality. Some of the early prediction methods used comparative approaches by homology mapping, see, e.g., [27, 43]. With the introduction of large gene database such as DEG, CEG and OGEE [4, 25, 40], researchers designed more complex prediction models using a wider set of features. These features can be broadly categorized into (i) sequence features, i.e., codon frequency, GC content, gene length [29, 35, 42], (ii) topological features, i.e., degree centrality, cluster coefficient [1, 6, 24, 31], and (iii) functional features, i.e., homology, gene expression cellular localization, functional domain and molecular properties [5, 9, 23, 30, 39].

Sequence based features can be directly obtained from the primary DNA sequence of a gene and its corresponding protein sequence. Functional features such as network topology requires knowledge of protein-protein interaction network, e.g., STRING and HumanNET [15, 37]. Gene expression and functional domain information can be obtained from databases like PROSITE and PFAM [10, 14]. Some of the less studied bacterial species, however, lack these functional and topological features, which prevents the use of classifiers that rely on them. Sequence based classifiers are the most practical methods because they use the minimal amount of features.

Several studies have been published on the problem of predicting essential genes from their sequence. In [35], the authors developed a tool called ZUPLS that uses (i) a Z-curve derived from the sequence, (ii) homology mapping and (iii) domain enrichment score as features to predict essential genes in twelve prokaryotes after training the model on two bacteria. Although ZUPLS worked well on cross-organism prediction, the limited number of bacterial species used as training dataset cast doubts on the ability of ZUPLS to generalize to more diverse bacterial species. In [22], the authors proposed a computational method that employs PCA on features derived from the gene sequence, protein domains, homologous and topological information. Among the studies that predicts essential genes across multiple bacterial species, [30] employed several genomic, physio-chemical and subcellular localization features to predict gene essentiality across fourteen bacterial species. In their work, the authors dealt with the redundancy in the dataset (i.e., homologous genes shared by multiple bacterial genomes) by clustering genes based on their sequence similarity. In [29], nucleotide, di-nucleotide, codon, and amino acid frequencies and codon usage analysis were used for predicting essentiality in sixteen bacterial species. The authors used CD-HIT [20] for homology detection in both essential and non-essential genes. In [28], the authors identified essential genes in fifteen bacterial species using information theoretical features, e.g., Kullback-Leibler divergence between the distribution of *k*-mers (*k* = 1, 2, 3), conditional mutual information and entropy features. Although their work showed promising results for intra-organism and cross-organism predictions, the model performed rather poorly when trained on the complete bacterial dataset. Recently, [23] showed the most extensive prediction analysis on thirty-one bacterial species. The authors employed the features proposed in [30], with additional features such as trans-membrane helices and Hurst exponent. Their algorithm used a regularized feature selection method called least absolute shrinkage and selection operator (Lasso) and used SVM as the classifier.

The latest work in gene essentiality prediction [2] uses network based features and Lasso for feature selection with Random Forest as classifier. The authors used a recursive feature extraction technique to compute 267 features in three different categories i.e. *local features* such as degree, *egonet features* which refers to the node and the induced subgraph formed by a node and all of its neighbors and *regional features* which is a combination of local and egonet features. They also used fourteen network centrality measures as a separate feature set for the essentiality prediction. Finally they combined their network based features with the sequence based features in [23] and [35] for their prediction model. For the models in [23], [2] and [35], the authors down-sampled non-essential genes to balance the training set but did not realize that their dataset contained multiple copies of homologous genes which created a “data leak” issue which biased their results (see below).

In this work we propose a feed forward deep neural network (DNN) called DeeplyEssential that uses features derived solely from the primary gene sequence to identify essential genes in bacterial species, thus maximizing its practical application compared to other predictors that require structural or topological features which might not be readily available. To the best of our knowledge, this is the first time a deep neural network has been used for gene essentiality prediction.

## 2 Materials and Methods

### 2.1 Dataset

Genetic data for thirty bacterial species were obtained from the database DEG, which is a curated and comprehensive repository of experimentally-determined bacterial and archaeal essential genes. Among the thirty bacterial species, nine are *Gram-positive* (GP) and twenty-one are *Gram-negative* (GN). DEG provides the primary DNA sequence and corresponding protein sequence for both essential and non-essential genes, as well as gene functional annotations. We only considered protein-coding genes, i.e., we excluded RNA genes, pseudo-genes and other non-coding genes. At the time of writing, DEG contained exactly 28,876 essential protein-coding genes (of which 8,746 belonged to a GP species and 20,130 belonged to a GN species) and 209,026 non-essential protein-coding genes (of which 45,002 were GP and 164,024 were GN). Table 1 shows the basic statistics of the dataset. Observe that the dataset is highly unbalanced: while species NC 000907 and NC 002771 have approximately the same number of essential and non-essential genes and bacteria NC 000908 has more essential genes than non-essential genes, for ten bacterial species less than 10% of their genes are essential. In order to improve the performance of our classifier, we balanced the dataset by downsampling non-essential genes.

**Table 1.** The thirty bacterial species used for our experiments (GP is Gram-positive, GN is Gram-negative)

| Accession | GP/GN | # Essential genes | # Non-essential genes |
| --- | --- | --- | --- |
| NC.000907 | GN | 1284 | 1024 |
| NC.000908 | GP | 762 | 188 |
| NC.000913 | GN | 1810 | 14000 |
| NC.000915 | GN | 646 | 2270 |
| NC.000962 | GP | 4144 | 17586 |
| NC.000964 | GP | 542 | 7808 |
| NC.002163 | GN | 788 | 5602 |
| NC.002505/002506 | GN | 1558 | 5886 |
| NC.002516 | GN | 906 | 21266 |
| NC.002745 | GP | 604 | 4562 |
| NC.002771 | GP | 620 | 644 |
| NC.003197 | GN | 460 | 8456 |
| NC.004347 | GN | 804 | 2206 |
| NC.004631 | GN | 1422 | 15822 |
| NC.004663 | GN | 650 | 8906 |
| NC.005966 | GN | 998 | 5188 |
| NC.006351/006350 | GN | 1010 | 10444 |
| NC.007297 | GP | 454 | 2674 |
| NC.007795 | GP | 702 | 5082 |
| NC.008463 | GN | 670 | 1920 |
| NC.008601 | GN | 784 | 2658 |
| NC.009009 | GP | 436 | 4104 |
| NC.009511 | GN | 1070 | 8630 |
| NC.010729 | GN | 1488 | 6870 |
| NC.011375 | GP | 482 | 2354 |
| NC.011916 | GN | 960 | 6448 |
| NC.016776 | GN | 1094 | 7486 |
| NC.016810 | GN | 706 | 8070 |
| NC.016856 | GN | 210 | 10420 |
| NC.007650/007651 | GN | 812 | 10452 |

### 2.2 Feature selection

As said, various intrinsic gene features, such as protein domains, protein-interaction network data, etc. have been used for predicting gene essentiality [22, 28]. DeeplyEssential utilizes codon frequency, maximum relative synonymous codon usage (RCSU), codon adaptation index (CAI), gene length and GC content. Along with these DNA-derived features, DeeplyEssential also uses amino acid frequency and sequence length from the protein sequences.

#### 2.2.1 Codon frequency

Codon frequency has been recognized an important feature for gene essentiality prediction [23, 30]. Given the primary DNA sequence of a gene, its codon frequency is computed by sliding a window of three nucleotides along the gene. The raw count of 4^3^ = 64 codons is then normalized by the total number of genes. Observe in Figure 1 that the codon frequency can be quite different in the two classes. For instance, codon AAA, GAA, TGA, GAT, AAG, ATT and AGA had at least 30% difference in their normalized codon frequency between essential and non-essential genes.

**Figure 1.**
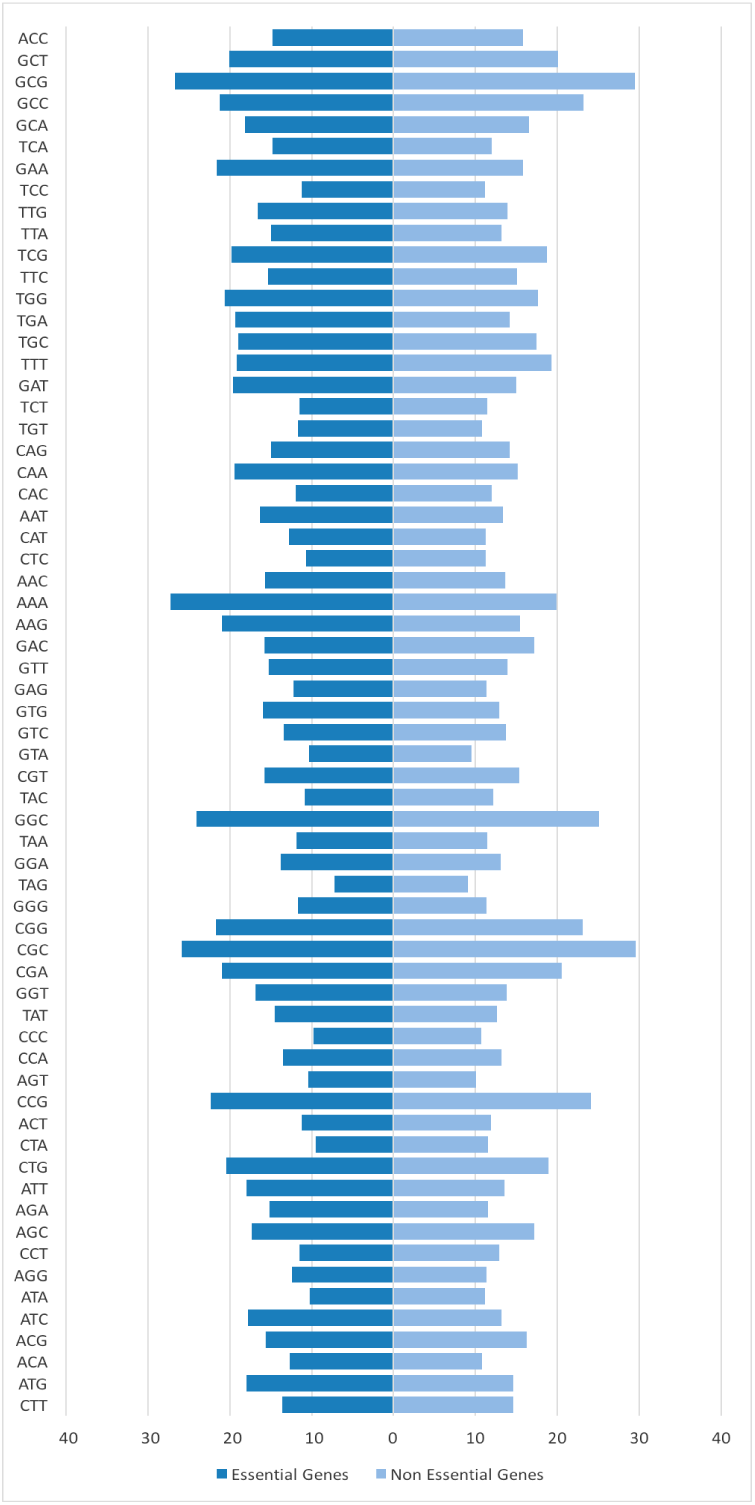
Normalized codon frequency of gene sequences in GP + GN dataset

#### 2.2.2 Gene length and GC content

Other distinguishing features for gene essentiality are gene length and GC content. Figure 2 shows the distribution of gene length in GP, GN and complete dataset (GP+GN). Observe that in the complete dataset and the GN dataset, gene have similar average length in the two classes, while in the GP dataset essential genes are on average longer than non-essential genes. As said, the GC content is another informative feature of essentiality prediction. Figure 3 shows the difference in distribution in GC content between two classes. Observe that non-essential genes have higher GC content than essential genes.

**Figure 2.**
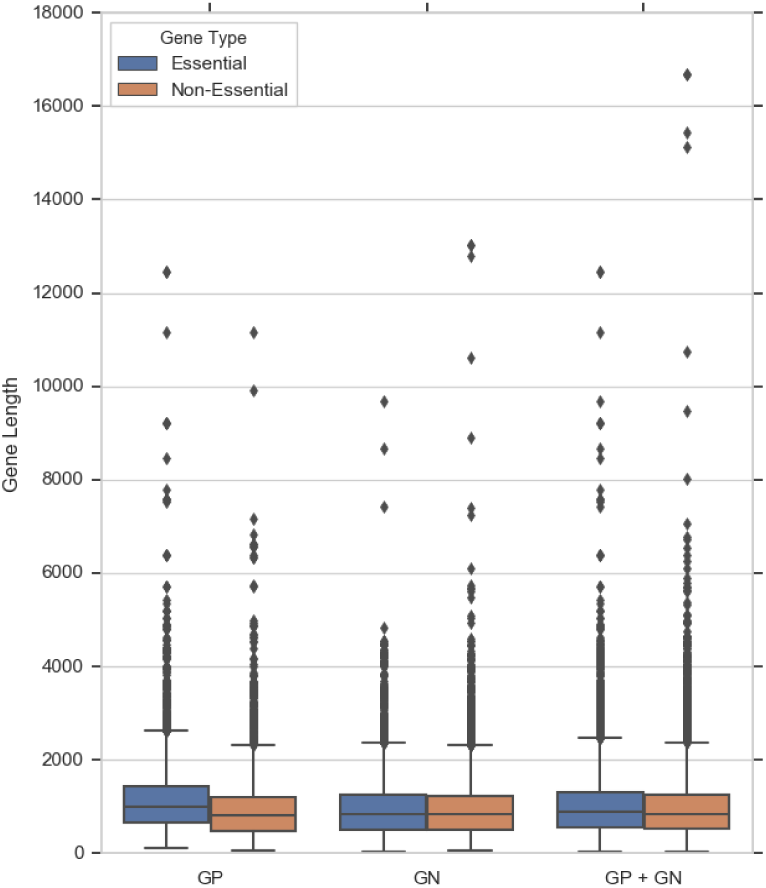
Distribution of gene lengths in datasets GP+GN, GN, GN

**Figure 3.**
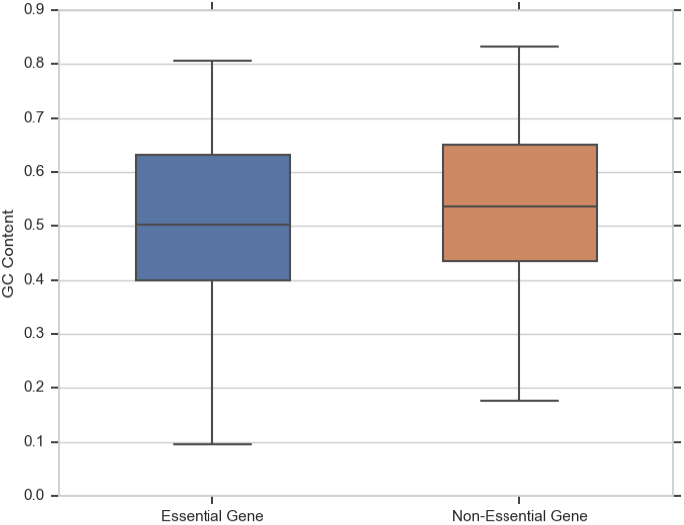
GC content distribution in essential and non-essential gene sets in GP + GN dataset

#### 2.2.3 Relative synonymous codon usage

Unbalanced synonymous codon usage is prevalent both in prokaryotes and eukaryotes [26]. The degree of bias varies among genes not only in different species but also among genes in the same species. Differences in codon usage in one gene compared to its surrounding genes may imply its foreign origin, different functional constraints or a different regional mutation. As a result, examining codon usage helps to detect changes in evolutionary forces between genomes. Essential genes are critical for the survival of an organism thus codon usage acts as a strong distinguishing feature. To calculate the relative synonymous codon usage we compare the observed number of occurrence of each codon to the expected number of occurrences (assuming that all synonymous codons have equal probability). Given a synonymous codon *i* that has an *n*-fold degenerate amino acid, we compute the *relative synonymous codon usage* (RCSU) as follows

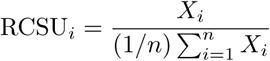

where *X*_*i*_ is the number of occurrence of codon *i*, and *n* is 1, 2, 3, 4, or 6 (according to the genetic code).

#### 2.2.4 Codon adaptation index

The *codon adaptation index* (CAI) estimates the bias towards certain codon that are more common in highly expressed genes [26]. The CAI is defined by the geometric mean of the relative adaptedness statistics. The *relative adaptedness* for codon *i* is defined on the relative frequency of the codon in a species-specific reference set of highly expressed genes. Formally, the relative adaptedness is defined by

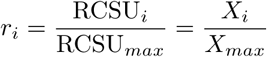

where RCSU_*max*_ and *X*_*max*_ are corresponding RCSU and *X* value of the most frequently used codon. The CAI for a gene is defined by

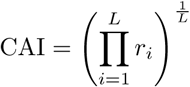

where *L* is the number of codons in the gene excluding methionine, tryptophan, and stop codon. The value of CAI ranges from zero to one, where zero indicates no bias.

#### 2.2.5 Protein sequence features

Another informative set of features used for the prediction of gene essentiality are those derived from the corresponding protein sequences. Previous studies have used frequency of rare amino acids, and the number of codons that are one-third base mutations removed from the stop codons [23]. DeeplyEssential only uses amino acids frequencies and the lengths of the protein sequences.

#### 2.2.6 Combining all the features

Given the primary DNA sequence of a gene, we generate 4^3^ = 64 values for the codon frequency, and one value for the GC content, gene length, CAI and RCSU_max_. From the protein sequence, we compute the amino acid frequency vector (20 components), and one value for the protein length. The total number of features used by DeeplyEssential is 89.

### 2.3 Multi-layer perceptron

Multi-layer perceptron (MLP) consists of multiple layers of computational units where the information flows in forward direction, from input nodes through hidden nodes to the output nodes without any cycles [33]. MLP networks have been used successfully for several molecular biology problems, see, e.g. [11, 21, 34]. The architecture of DeeplyEssential is composed of an input layer, multiple hidden layers and an output layer. The output layer encodes the probability of a gene to be essential. The addition of dropout layer makes the network less sensitive to noise in the training and increase its ability to generalize. This layer randomly assign zero weights to a fraction of the neurons in the network [36].

Let 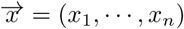 be the input to the MLP. Let vector *y* denotes the output of the *i*^*th*^ hidden layer. The output *y* depends on the input in the previous layer as follows

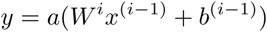

where *a* is the activation function, *b* is the bias and *W* is the weight matrix for each edge in the network. During training, the network learns the weights *W* and the bias *b*. DeeplyEssential uses a rectified linear unit (ReLU) in each neuron in the hidden layers. ReLU is an element-wise operation that clamps all negative values to zero.

In the output layer DeeplyEssential uses a sigmoid as the activation function to perform discrete classification

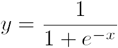

The loss function is binary cross-entropy defined by

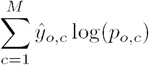

where *M* is the number of classes (two in our case), *ŷ* is the binary indicator if class label *c* is the correct classification for observation *o*, and *p* is the predicted probability observation *o* is of class *c*. Figure 4 illustrates the architecture of the neural network used in DeeplyEssential.

**Figure 4.**
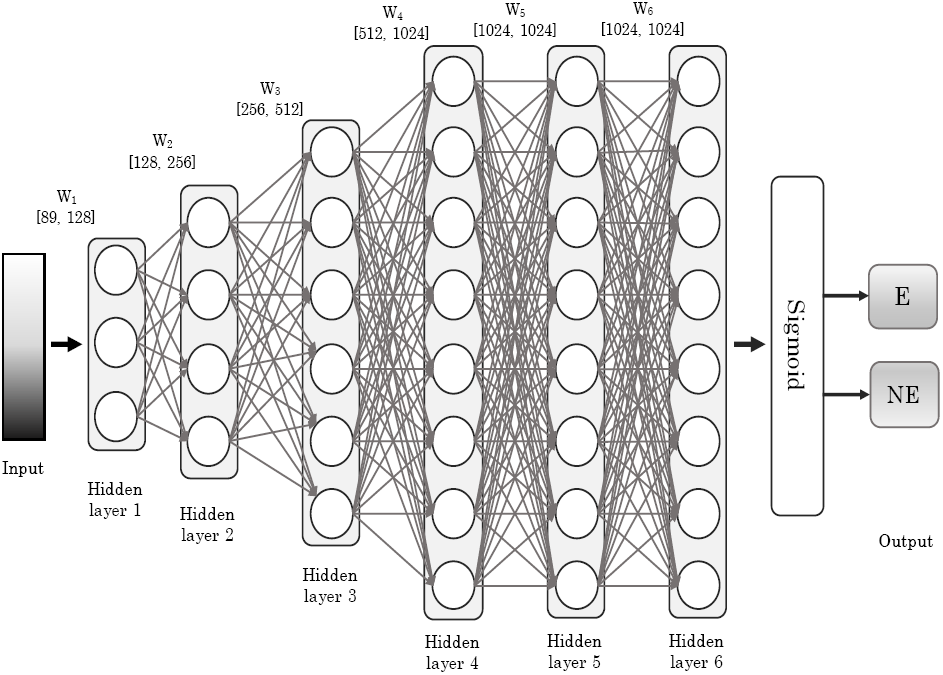
The architecture of the neural network used in DeeplyEssential

## 3 Results and Discussion

### 3.1 Classifier design and evaluation

As mentioned in Section 2.1, the number of non-essential genes is significantly larger than the number of essential genes. To address this imbalance in the training set and allow for unbiased learning, we randomly down-sample non-essential genes. In [42], the authors showed that balancing the dataset did not negatively influence the prediction of gene essentiality.

#### 3.1.1 Model hyper-parameters

Recall that each gene (and its corresponding protein) is represented by 89 features in the input layer. The deep learning architecture of DeeplyEssential was determined my running extensive experiments on the training data over a wide range of hyper-parameters. The number of hidden layers, the number of nodes in each of the hidden layers, the batch size, the dropout rate and the type of optimizer were selected by optimizing the performance of the classifier. Table 2 lists the range of hyper-parameter considered and the values of the hyper-parameter selected for the final architecture of DeeplyEssential.

**Table 2.** Hyperparameters for DeeplyEssential

| Parameters | Range | Selected Parameter |
| --- | --- | --- |
| # hidden layers | [2 - 8] | 6 |
| # nodes | [32, 64, 128, 512, 1024, 2048] | [128, 256, 512, 1024, 1024, 1024] |
| dropout rate | [0.1 - 0.5] | 0.3 |
| epochs | — | 100 (early stopping) |
| optimizer | sgd, adam, adadelata, RMSProp | adadelata |

Observe in Figure 4 that the final fully-connected layer reduces the 1024 dimensional vector to a two-dimensional vector corresponding to the two prediction classes (essential/non-essential). The sigmoid activation function forces the output of the two neurons in the output layer to sum to one. Thus their output value represents the probability of each class. Among the available optimizer in Table 2, we chose adadelta because of its superior performance. Adadelta is parameter-free, thus we do not need to define the learning rate. The training was ran for 100 epochs with early stopping criteria.

We trained DeeplyEssential on three datasets, namely GP, GN and GP+GN (see Section 2.1 and Section 3.2). For each dataset, 80% data is used for training, 10% data for validation and 10% data for testing. The random selection was repeated ten times, i.e., a ten-fold cross-validation was performed to complete the inference.

#### 3.1.2 Evaluation metrics

The tools described in [23], [30], [29] and [28] are currently unavailable. We ran DeeplyEssential on the datasets used in the corresponding papers, and compared DeeplyEssential’s classification metrics to the published metrics.

We evaluated the performance of DeeplyEssential using the Area Under the Curve (AUC) of the Receiver Operating characteristic Curve (ROC). ROC plot represents the trade-off between sensitivity and specificity for all possible thresholds. Although our primary evaluation measure is the AUC score, we report the following additional performance measures

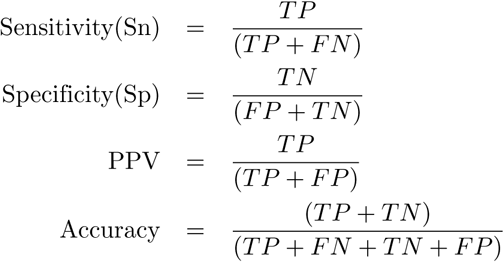

where *TP, TN, FP* and *FN* represent the number of true positives, true negatives, false positives and false negatives, respectively.

All experiments were carried out a Titan GTX 1080 Ti GPU, running Keras 2.1.5.

### 3.2 Gene essentiality prediction

We collected essential and non-essential gene for thirty bacterial species as described in Section 2.1 into three datasets, namely GP, GN and GP+GN. After re-balancing the dataset by down-sampling non-essential genes, we extracted the features for each gene as explained in Section 2.2. Table 3 shows the basic statistics for each dataset.

**Table 3.** Basic statistics for GP, GN and GP+GN (balanced and unbalanced)

| Dataset | # Training Samples | # Validation Samples | # Test Samples |
| --- | --- | --- | --- |
| GP | 7,065 | 883 | 884 |
| GN | 14,364 | 1,795 | 1,797 |
| GP+GN (bal) | 21,432 | 2,678 | 2,680 |
| GP+GN (unbal) | 90,571 | 11,321 | 11,322 |

Table 4 shows the training classification performance of DeeplyEssential, averaged over ten repetitions. The violin plot in Figure 5 shows the distribution of AUCs across the ten repetitions of the experiment, which appears very stable. The receiver operator curves (ROC) are shown in Figure 7. DeeplyEssential yielded an area under the curve of 0.838, 0.829 and 0.842 for GP, GN and GP+GN on average, respectively. The ROC curve also indicates the relation between the number of training samples and stability in model performance. Observe that DeeplyEssential’s performance was more stable on the GP+GN dataset than the GP dataset (which contains the smallest number of samples).

**Table 4.** Training classification performance of DeeplyEssential on GP, GN, GP+GN

| Metric | GP | GN | GP+GN |
| --- | --- | --- | --- |
| AUC | 0.838 | 0.823 | 0.842 |
| Sensitivity | 0.741 | 0.784 | 0.801 |
| Specificity | 0.758 | 0.708 | 0.721 |
| PPV | 0.774 | 0.722 | 0.749 |
| Accuracy | 0.749 | 0.745 | 0.762 |

**Figure 5.**
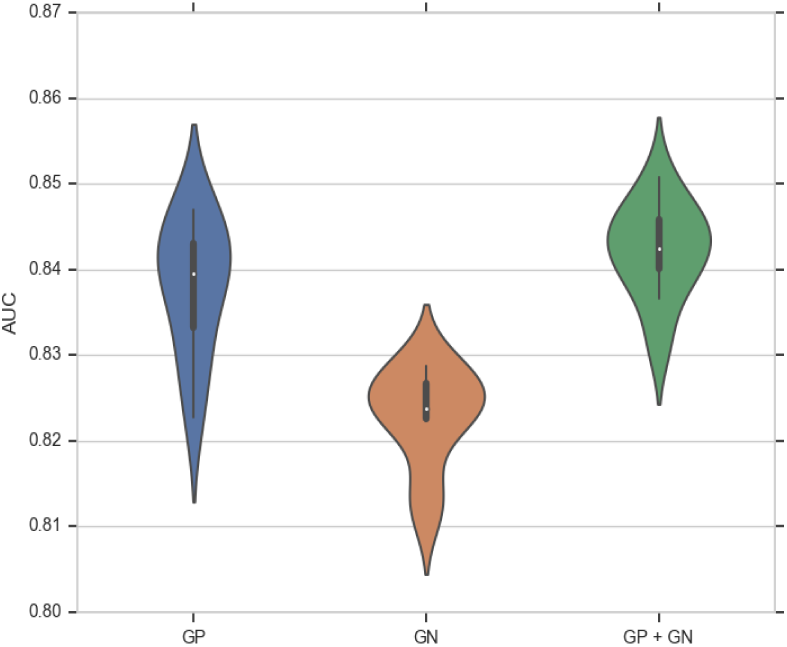
Violin plot of DeeplyEssential’s AUC across ten experiments on the GP, GN, GP+GN datasets

**Figure 6.**
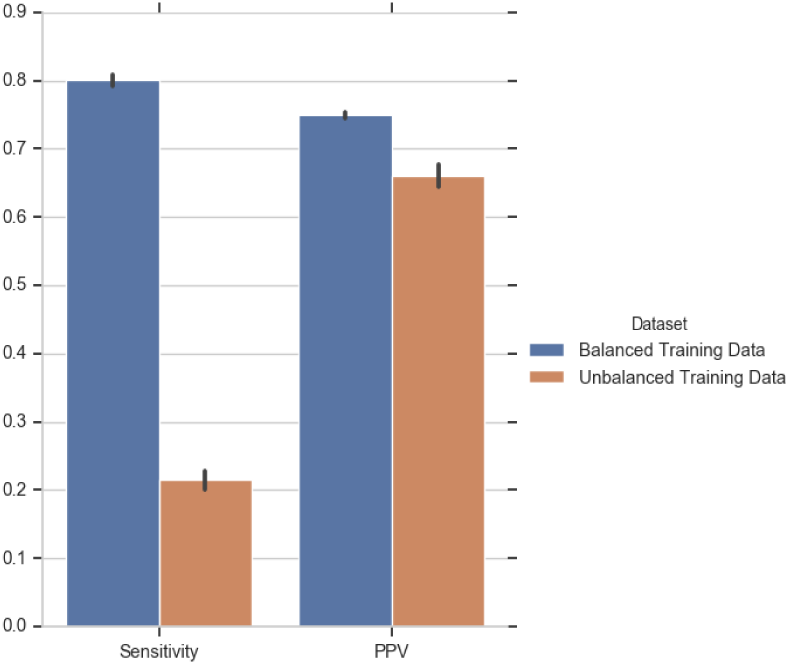
Comparing the prediction performance of DeeplyEssential when trained on balanced or unbalanced GP+GN dataset

**Figure 7.**
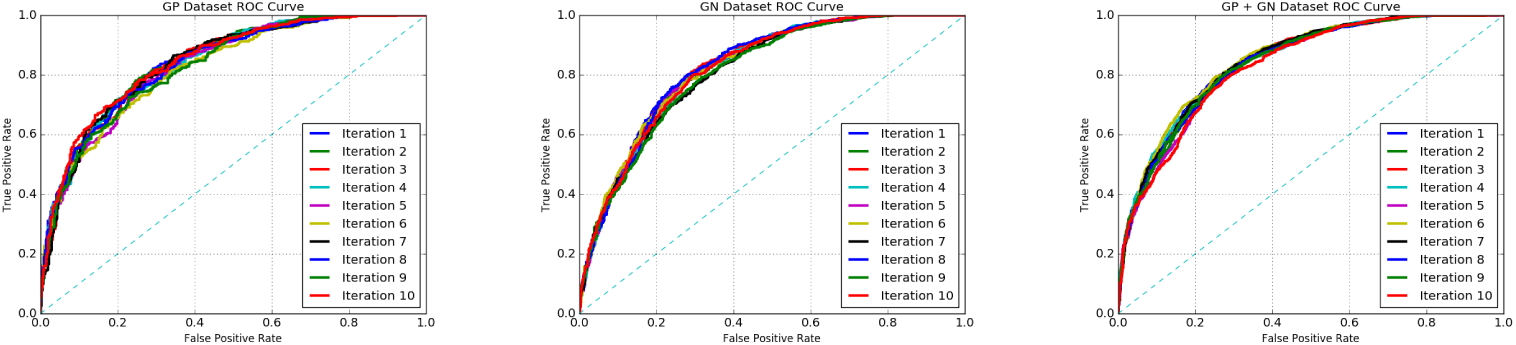
DeeplyEssential’s ROC curves on GP, GN, GP+GN

### 3.3 Comparison with published methods that use down-sampling

As said in Section 2.1, the gene essentiality dataset is highly unbalanced. It is well-known that class imbalance can negatively affect the performance of a classifier [41]. To quantify how class imbalance affects the performance of our classifier we trained DeeplyEssential on the full (unbalanced) dataset that has 322.6% more non-essential genes than essential genes. Figure 6 shows that the sensitivity and Positive Predictive Value (PPV) of the classifier trained on unbalanced data is much worse than the balanced dataset. As said, some of the existing methods use down-sampling to address this problem. Both Liu *et al*. 2017 [23] and Azhagesan *et al*. 2018 [2] randomly down-sampled the majority class data to match the size of the minority class. DeeplyEssential also uses this approach. Table 5 shows the performance

**Table 5.**
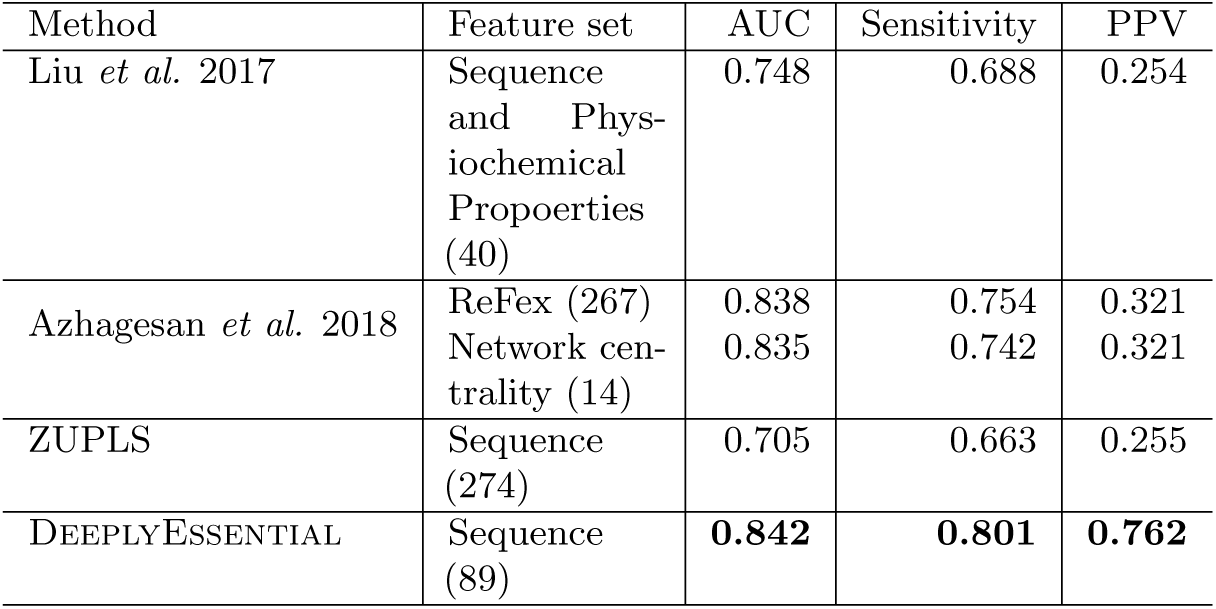
Comparing the performance of DeeplyEssential on down-sampled dataset; numbers in boldface indicate the best performance

| Method | Feature set | AUC | Sensitivity | PPV |
| --- | --- | --- | --- | --- |
| Liu *et al.* 2017 | Sequence and Physiochemical Properties (40) | 0.748 | 0.688 | 0.254 |
| Azhagesan *et al.* 2018 | ReFex (267) | 0.838 | 0.754 | 0.321 |
|  | Network centrality (14) | 0.835 | 0.742 | 0.321 |
| ZUPLS | Sequence (274) | 0.705 | 0.663 | 0.255 |
| DEEPLYESSENTIAL | Sequence (89) | **0.842** | **0.801** | **0.762** |

DeeplyEssential compared to the two published methods that use down-sampling. Observe that DeeplyEssential achieves the best AUC, sensitivity and PPV.

### 3.4 Identification of “data leak” in the gene essentiality prediction

Bacteria are unicellular organisms with a relatively small set of genes. Across bacterial species a significant fraction of genes are conserved because they performs similar fundamental biological functions. These conserved (homologous) genes are quite similar at the sequence level. All published methods rely on dataset containing multiple bacteria on which genes have been labeled essential or non-essential. Let *x* and *y* be two homologous genes, i.e., *x* and *y* have very similar sequence. If *x* is used on the training and *y* if used for testing, this introduces a bias, or a “data leak”. Training examples and testing examples are supposed to be distinct, and in this hypothetical scenario they are not.

To quantify the effect of the data leak issue, we clustered the set of all genes across the thirty bacterial species using OrthoMCL [19]. OrthoMCL is a popular method for clustering orthologus, homologous and paralog proteins which uses reciprocal best hit alignment to detects potential in-paralog/recent paralog pair, and reciprocal alignments best hits across any two genomes to identify potential ortholog pairs. A similarity graph is then generated based on the proteins that are interlinked. To split large clusters, a Markov Clustering algorithm (MCL) is then invoked [38]. Inside MCL clusters, weights between each pair of proteins is normalized to correct for evolutionary differences.

As said, OrthoMCL produces a list of clusters where each cluster consists of genes that have been determined to be orthologus. To quantify the effect of gene sequence similarity on the prediction performance, we created a dataset where no gene from a single cluster can end up in both the training set and the testing set. The modified dataset contains 11,168 training samples, 2,798 validation samples and 4,270 testing samples. The prediction was repeated ten times. Table 6 shows the clustering step heavily influences DeeplyEssential’s prediction performance: AUC decreased by more than 7% (on average), while the accuracy decreased by 6.9% (along with significant decrease in all performance measures). Figure 8 shows the difference in performance before and after clustering. While the AUCs were stable across experiments, sensitivity, specificity and PPV varied largely across experiments for clustered dataset.

**Table 6.** Comparing the effect of clustering on the prediction performance of DeeplyEssential on the GP+GN dataset

| Metric | Non-clustered | Clustered | Difference (%) |
| --- | --- | --- | --- |
| AUC | 0.842 | 0.786 | 7.12% |
| Sensitivity | 0.801 | 0.780 | 2.69% |
| Specificity | 0.721 | 0.646 | 11.60% |
| PPV | 0.749 | 0.688 | 8.86% |
| Accuracy | 0.762 | 0.713 | 6.87% |

**Figure 8.**
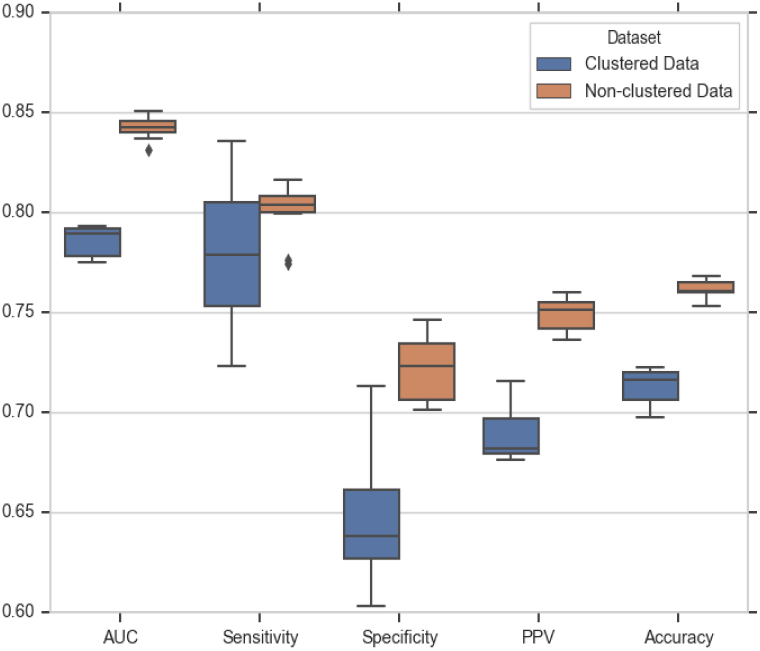
Effect of “data leak” on DeeplyEssential’s prediction performance

### 3.5 Comparison with methods that address orthologus genes

Some published studies have addressed the data leak issue by identifying homologus genes using sequence similarity metrics. In [28], the authors used the Kullback-Leibler divergence (KLD) to measure the distance between *k*-mer distribution (for *k* = 1, 2, 3) obtained from sequences. In [29], the authors used CD-HIT to remove redundancy in the training data and improve the generalization ability of their model. As explained in the previous section, DeeplyEssential uses OrthoMCL to cluster homologous genes to prevent similar genes to appear in both training and testing dataset. Table 7 and Table 8 shows the performance comparison of DeeplyEssential with [29] and [28] on their respective datasets. Observe that in both cases DeeplyEssential achieves the best predictive performance.

**Table 7.** Comparing the performance of DeeplyEssential and Ning *et al* on the Ning *et al* dataset [29]; numbers in boldface indicate the best performance

| Method | Clustering method | AUC |
| --- | --- | --- |
| Ning *et al* 2014 | CD-HIT | 0.758 |
| DEEPLYESSENTIAL | OrthoMCL | **0.818** |

**Table 8.**
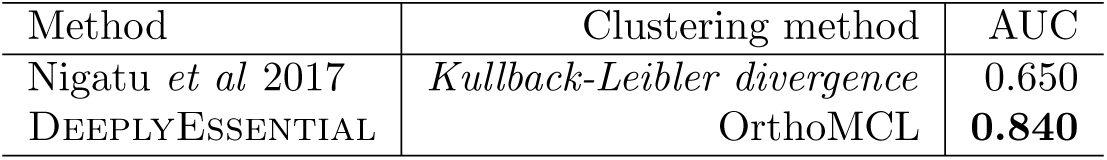
Comparing the performance of DeeplyEssential and Nigatu *et al* on the Nigatu *et al* dataset [28]; numbers in boldface indicate the best performance

### 3.6 Feature importance

DeeplyEssential uses exclusively sequence based features and yet produces higher prediction performance. Unlike other machine learning classifiers, the DNN architecture does not readily provide any insight about the feature set that contributed maximally towards the prediction performance. To understand the impact of a feature on the predictive performance, we carried out an ablation study which removes feature(s) from the input and observe the performance difference to determine the importance of a feature. However, this type of study is not very informative in the presence of highly correlated features. In this case, the absence of a feature can be compensated by another feature which is highly correlated with the former feature. To address this issue, we first computed pairwise Pearson correlation among all input features. Figure 9 illustrate the heatmap of the pairwise correlation. Each axis shows the indices of the features: indexes 0–65 contains DNA specific feature, index 68–89 contains protein specific features. GC content, CAI and *RSCU* _*max*_ have negative correlation with all other features. There were nineteen pair of features showing correlation higher than 0.9 (in absolute value). For our ablation study we either removed one feature at a time (if uncorrelated) or one of the 19 feature pairs to test the performance changes on the GP + GN dataset using 5-fold cross validation. We measured the difference in AUC and ordered the features based on their impact in decreasing the predictive performance (Figure 10). Observe that codon TTT caused the highest AUC decrease (3.5%) while AGA, TTC, CGT, CGA, gene length, protein length, GC content, CAI, amino acids R, W, Y, K, L and pairs of correlated features CCG+CGC, TAA+TTA, Gene length+L, D, and protein length+T caused more than a 3% AUC decrease.

**Figure 9.**
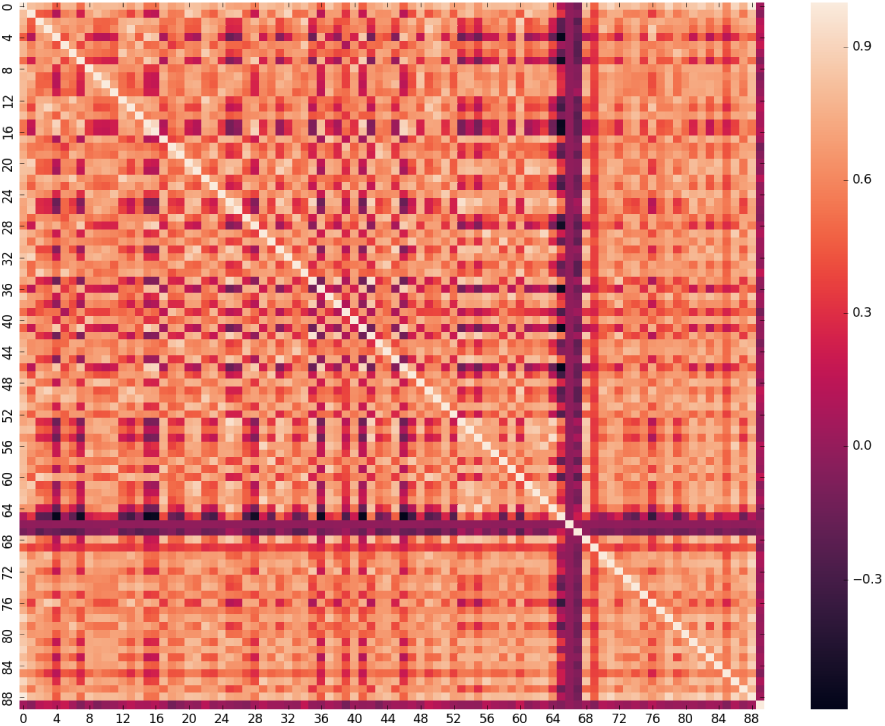
Pairwise correlation among all features; features 0–65 are DNA specific feature; features 68–89 are protein specific features

**Figure 10.**
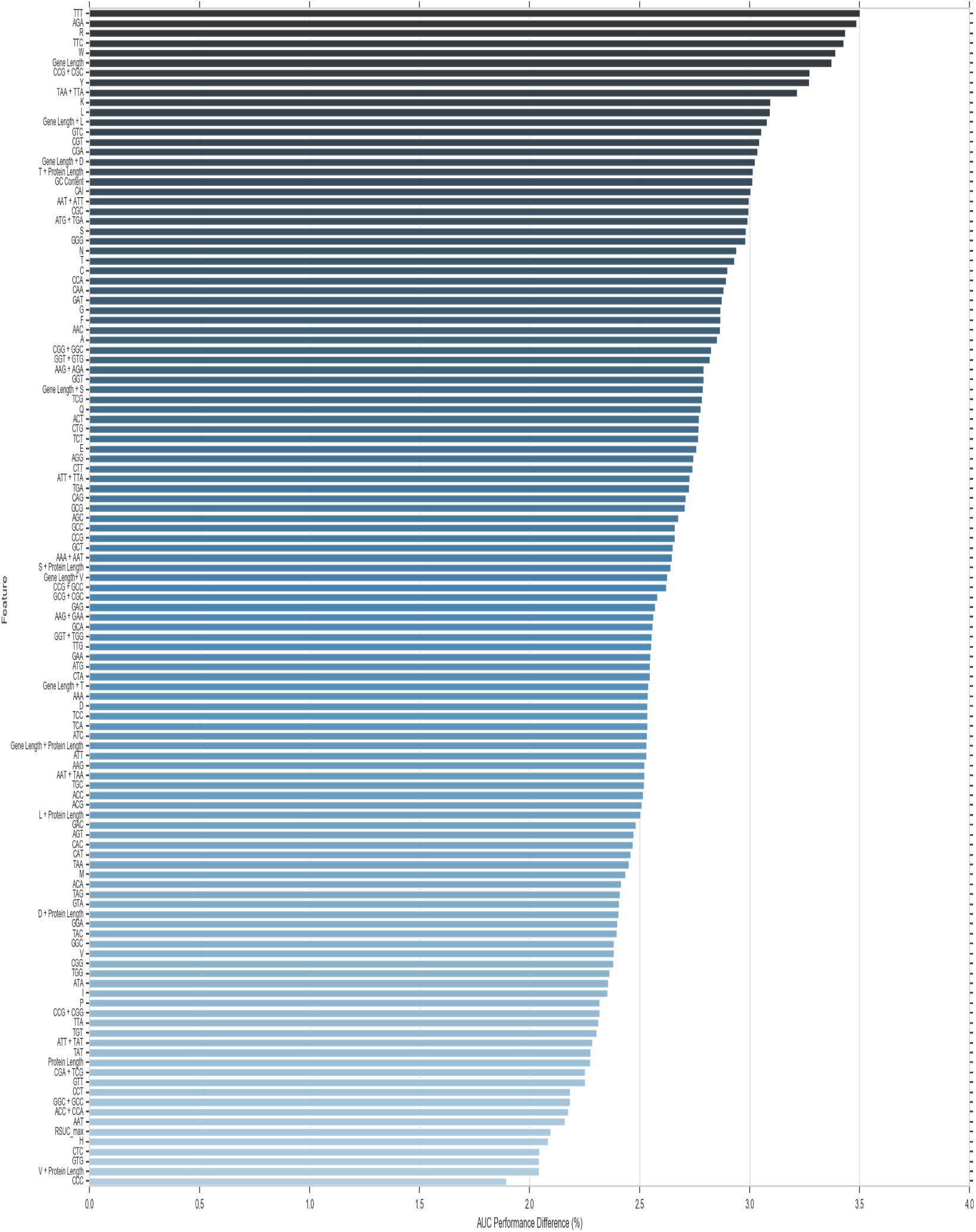
Changes in AUC predictive performance due to the removal of a feature or pairs of correlated features

### 3.7 Discussion

A large number of structural and functional features have been used for gene essentiality prediction, i.e. producibility, choke points, load scores, damages, degree of centrality, clustering coefficient, closeness centrality, betweenness centrality, gene expression, phyletic retention, among others. These features cannot be obtained from the gene sequences and are often not available for many bacterial species. To maximize its practical utility, DeeplyEssential uses exclusively features derived directly from the sequence.

Our experiments showed that DeeplyEssential has better predictive performance both on down-sampled and clustered datasets. On the down-sampled dataset used in [23], DeeplyEssential showed an improvement of 12.8% in AUC compared to [23] and achieved a slightly better AUC on the network-based feature model [2]. In addition, DeeplyEssential produced significantly better sensitivity and precision than the three methods in Table 5, achieving 6.2% improved sensitivity and 137.4% improved precision compare to [2]. If one uses all the 597 features in the prediction model in [2], then this latter method achieves 1.7% improved AUC compared to DeeplyEssential. We believe that collecting this very large amount of features from multiple databases does not warrant the additional (minor) benefit in predictive performance. DeeplyEssential also achieved better performance on clustered datasets. Table 7 and Table 8 show 7.9% and 29.2% improved AUC compared to [29] and [28], respectively.

## 4 Conclusion

We proposed a deep neural network architecture called DeeplyEssential to predict gene essentiality in microbes. DeeplyEssential makes minimal assumption about the input data (i.e, it only uses the gene sequence), thus maximizing its practical application compared to other predictors that require structural or topological features which might not be readily available. Extensive experiments shows that DeeplyEssential has better predictive performance than existing prediction tools. We believe that DeeplyEssential could be further improved if more annotated bacterial data was available, making it an essential tool for drug discovery and synthetic biology experiments in microbes.

